# Trans-ancestry Mendelian Randomization Discovers Novel Causal Genes for Autoimmune Disease Traits

**DOI:** 10.1101/2025.04.17.649410

**Authors:** Siyuan Chen, Chen Wang, Kent D. Taylor, Jerome I. Rotter, Xiuqing Guo, Stephen S. Rich, Ani Manichaikul, Dajiang Liu

## Abstract

Mendelian randomization (MR) using summary statistics from genome-wide association studies (GWAS) has become a powerful tool for dissecting the causal relationships between exposure and outcomes. As GWAS begins to incorporate samples of diverse ancestries, the optimal strategy to perform MR analysis integrating these GWAS results remains to be determined. To fill in this gap, we proposed a robust method that aggregates the genetic variant–risk factor association summary statistics of multiple ancestries by modelling the genetic effects of the exposure as a function of the principal components of genome-wide allele frequency. This allows borrowing information from different ancestry groups and accounting for potential between-ancestry heterogeneities. The method is general and can be used with different MR methods. We demonstrated that our method improves accuracy and power compared to the inverse-variance weighting method based on only ancestry-stratified samples, with additional benefits in correcting winner’s curse bias. We also illustrated the flexibility and potential of discovering novel causal genes for autoimmune disease traits of our method in practice.

## Introduction

Mendelian Randomization (MR) has been a popular tool for inferring causal relationships between factors of interest (exposure) and outcomes in observational studies. With genetic variants (e.g., SNPs) as instrumental variables (IVs), the inference of causal effects can be made for the exposures that are heritable, using GWAS summary statistics as input. In two-sample MR, the variant-exposure and variant-outcome associations are derived from two non-overlapping samples, and ideally, they should be from the same underlying population[1]. When multi-ancestry GWAS summary statistics are available, standard inverse variance weighted (IVW) MR analysis would need to be conducted in each ancestry separately, which directly leads to a reduction in effective sample size, and hence possibly depletion in performance.

The genetic effect may differ between ancestries. More than simply pooling multiple ancestry cohorts together using fixed-effect meta-analysis may be underpowered. Previously, we developed methods that model the genetic effects of each cohort as a function of the principal components of genome-wide allele frequencies [2]. This adjustment allows us to accommodate the heterogeneities of genetic effect sizes between ancestries and borrow strength across ancestries to improve the precision of the genetic effect estimates. Extending the framework to improve multi-ancestry MR would be necessary to improve the power of MR methods.

Furthermore, a two-sample setting raises concern for the winner’s curse bias, which occurs when the discovery GWAS for IV selection is also used for association estimation[3]. It is well known that genetic effects for variants selected as IVs tend to be inflated in the original discovery data set, especially when the power is low for genome-wide association studies. This may induce downward biases in the IVW estimator, and hence, a three-sample design is recommended, but it is not always feasible. Re-estimating the IV-exposure association using multi-ancestry summary statistics partially corrects for winner’s curse bias. It can be extended by incorporating the conditional likelihood method by Ghosh et al. [4] to include the conditional probability of the IV selection event within only matched global ancestry.

Here, we present our method, transMR, which aggregates information from multi-ancestry GWAS summary statistics for MR analysis. Compared to the IVW method, our model allows us to obtain a more plausible estimation of variant-exposure association from available summary statistics for the target population, i.e., of which the outcome cohort is. This is helpful especially when the sample size of the target population is small, as we will demonstrate in our simulation. In addition, our method brings adjustment on only associations and choice of instruments, which can be combined with other popular MR methods which focus on tackling issues like pleiotropic effects, such as GSMR[5]. Furthermore, we implement transMR in large-scale multi-ancestry studies that combine expression quantitative trait loci and GWAS. Our results identify more plausible causal traits in cohorts of relatively small sample sizes, as shown in our data application in causal gene discovery for autoimmune diseases.

## Results

### Method Overview

In a univariate MR analysis, our goal is to assess whether the exposure of interest has a causal effect on the outcome by utilizing genetic instrumental variables. Genetic variants serving as instruments must satisfy the three core IV assumptions, where the IVs should be associated with the exposure, uncorrelated with the confounders, and the IVs should only be associated with the outcome via their effects on the exposure (see Methods and Materials). The exchangeability assumption requires that instruments should not be confounded by confounders of the exposure-outcome relationship. At the same time, it is known that the distribution of SNPs and genetic effects can differ by ancestry structure. When only summary statistics are available, in a two-sample setting, the GWAS summary statistics should come from the same underlying population, otherwise the values of true IV-exposure/outcome effect may not be identical. Finally, the first-order inverse-variance weights are used to calculate the IVW estimate, and this assumes that the standard error of the estimated IV-exposure association is zero (the NOME assumption[6]).

We aim to find a better strategy for utilizing the GWAS results from multi-ancestry studies. Our method, transMR, uses a weighted meta-regression to model the genetic variant-exposure association (‘s) with principal components of the allele frequency (AF) matrix, which serves as quantitative proxies of ancestry. Estimation from each ancestry is weighted by the inverse of its variance in the regression. To avoid overfitting, we also included a ridge penalty on the regression parameters for the PCs as ancestry proxies. Such a model can be extended to further correct for winner’s curse bias brought by the two-sample design by replacing the likelihood contribution of estimation of *β* from the target population into a conditional likelihood that is conditional on the genetic effect. To avoid overfitting, we also included a ridge penalty on the regression parameters for the PCs as ancestry proxies. Such a model can be extended to further correct for winner’s curse bias brought by the two-sample design by replacing the likelihood contribution of estimation of *β* from the target population into a conditional likelihood that is conditional on the genetic effect being significant according to a pre-specified significance threshold (Details in Methods and Materials). The winner’s curse correction is performed for each SNP separately. Based on the proposed model, one can obtain an adjusted estimation of *β* in each population (possibly of different ancestries) on which the genetic variant-outcome association (Γ) was estimated. The improved genetic effect estimates can be used with any MR method.

Furthermore, when the target population for inference has a relatively small sample size (which is likely the case for a non-European population) and more significant uncertainty in estimated association arises, the violation of the NOME assumption is unignorable, transMR can borrow information from other population of a decent sample size to reduce such uncertainty and make a better estimation for the causal effect while still residing in the IVW framework.

### Trans-ancestry MR increases power for underrepresented population

We refer to the method without conditional likelihood transMR, while the one with correction for winner’s curse bias as transMR-c and that uses BIC to select the ridge penalty parameter as transMR- cBIC. We evaluate the performance of transMR methods through numerical simulations. In detail, we investigated the accuracy and precision of estimating the true causal effect θ = 0.5 with transMR and the original IVW method when all IVs are relatively strong and valid. We also considered two study designs. In the two-sample design, summary statistics for the exposure and outcome are obtained from two sets of independent samples, while in the three-sample design, another separate set of samples is used to select genetic variants with significant p-values as IV. Note that the three-sample design serves as a gold standard in our simulation. We also compared the power of proposed methods by setting true causal effect θ = 0.1 and the following extra two scenarios: (1) most of the IVs are weak; (2) the target population is underrepresented. The heterogeneity in genetic effects on the exposure is controlled by a scale parameter φ ∈ {0,0.5,1,1.5}. We start by generating individual-level data of multi-ancestry samples, running ancestry-stratified single-trait GWAS analysis, and then following a standard MR analysis workflow based only on summary-level data. We select IVs with a fixed p-value threshold and finally make inference on θ with candidate methods respectively.

We ran about 2000 iterations for each scenario for accuracy evaluation. In each case of SNP-exposure effect, along with no pleiotropy but unmeasured confounding, our transMR method proves to have a smaller bias, comparable variance and, in general, smaller MSE. The transMR-c and transMR-cBIC constantly have larger variances than other methods, but this also leads to higher coverage probability. Table (1) gives a detailed statistics table.

**Table 1.** Average bias, mean squared errors (MSE) and coverage probability (CP) for transMR and IVW methods.

| Methods | Bias | MSE | CP |
| --- | --- | --- | --- |
| $\varphi = 0$ | | | |
| transMR | -0.021 | 0.0035 | 0.88 |
| transMR-c | -0.028 | 0.0046 | 0.86 |
| transMR-cBIC | -0.024 | 0.0058 | 0.91 |
| IVW-2Sample | -0.033 | 0.0041 | 0.80 |
| IVW-3Sample | -0.012 | 0.0032 | 0.91 |
| $\varphi = 0.5$ | | | |
| transMR | -0.024 | 0.0036 | 0.86 |
| transMR-c | -0.031 | 0.0047 | 0.85 |
| transMR-cBIC | -0.026 | 0.006 | 0.9 |
| IVW-2Sample | -0.036 | 0.0043 | 0.78 |
| IVW-3Sample | -0.015 | 0.0032 | 0.91 |
| $\varphi = 1$ | | | |
| transMR | -0.024 | 0.0036 | 0.85 |
| transMR-c | -0.033 | 0.0052 | 0.87 |
| transMR-cBIC | -0.025 | 0.0068 | 0.9 |
| IVW-2Sample | -0.036 | 0.0043 | 0.79 |
| IVW-3Sample | -0.015 | 0.0032 | 0.91 |
| $\varphi = 1.5$ | | | |
| transMR | -0.022 | 0.0036 | 0.87 |
| transMR-c | -0.039 | 0.0067 | 0.82 |
| transMR-cBIC | -0.024 | 0.0086 | 0.92 |
| IVW-2Sample | -0.035 | 0.0041 | 0.80 |
| IVW-3Sample | -0.014 | 0.0031 | 0.91 |

We also investigated the power and type-I errors of the proposed methods with around 5000 iterations. All methods have controlled type-I error, and there is a slight improvement in power from transMR and transMR-c when (1) no requirement of IV is violated; (2) most SNP effects are relatively small. We want to point out that when the target population is of a small sample size, transMR remarkably improved in power (see Table (2)). In detail, IVW estimators are biased towards 0, although they still have a smaller variance. Notably, transMR methods outperformed the three-sample design with a relatively small population for discovery GWAS, suggesting that even though a separate cohort is available for GWAS and IV selection for underrepresented population, the performance is not as good as directly borrowing information from cohorts of a larger sample size but of different ancestry.

**Table 2.** Power of transMR methods and IVW methods with underrepresented population. *β*_j_ denotes the true genetic effects on the exposure.

| method | power |  |  |  |
| --- | --- | --- | --- | --- |
| non-European population proportion = 5%, $\beta_j \sim N(0.1, 0.05^2)$ | | | | |
| $\varphi$ | 0 | 0.5 | 1 | 1.5 |
| transMR | 0.9272 | 0.9252 | 0.9252 | 0.9268 |
| transMR-c | 0.9268 | 0.922 | 0.9232 | 0.9272 |
| IVW-2Sample | 0.7458 | 0.7442 | 0.7442 | 0.7458 |
| IVW-3Sample-small reference population | 0.8042 | 0.799 | 0.799 | 0.8042 |
| IVW-3Sample-large reference population | 0.9996 | 0.9996 | 0.9996 | 0.9996 |
| non-European population proportion = 10%, $\beta_j \sim N(0.08, 0.05^2)$ | | | | |
| transMR | 0.939 | 0.9362 | 0.9386 | 0.9406 |
| transMR-c | 0.9338 | 0.933 | 0.9334 | 0.935 |
| IVW-2Sample | 0.8222 | 0.8108 | 0.8146 | 0.8136 |
| IVW-3Sample-small reference population | 0.8502 | 0.8458 | 0.8472 | 0.8524 |
| IVW-3Sample-large reference population | 0.9914 | 0.99 | 0.99 | 0.9906 |
| non-European population proportion = 10%, $\beta_j \sim N(0.05, 0.05^2)$ | | | | |
| transMR | 0.487 | 0.4846 | 0.486 | 0.4892 |
| transMR-c | 0.4888 | 0.4848 | 0.4856 | 0.4852 |
| IVW-2Sample | 0.2978 | 0.3014 | 0.293 | 0.2978 |
| IVW-3Sample-small reference population | 0.338 | 0.3348 | 0.3382 | 0.3324 |
| IVW-3Sample-large reference population | 0.8746 | 0.8758 | 0.8656 | 0.8672 |
| non-European population proportion = 20%, $\beta_j \sim N(0.1, 0.05^2)$ | | | | |
| transMR | 0.999 | 0.999 | 0.999 | 0.999 |
| transMR-c | 0.999 | 0.998 | 0.999 | 0.998 |
| IVW-2Sample | 0.998 | 0.999 | 0.998 | 0.999 |
| IVW-3Sample-small reference population | 0.999 | 0.999 | 0.998 | 0.998 |
| IVW-3Sample-large reference population | 0.999 | 0.999 | 0.999 | 0.999 |

### Trans-ancestry MR discovers novel causal genes for autoimmune disease traits in East Asian population

Autoimmune diseases (AiDs), e.g., systemic lupus erythematosus, have a strong genetic influence[7]. Genome-wide association studies have identified hundreds of loci associated with several AiDs, but actual causal genes remain unclear, especially for the East Asian population, due to limited data availability.

We first conducted expression quantitative trait loci (eQTL) analysis on the gene expression of human peripheral blood mononuclear cells (PBMC) from the Multi-Ethnic Study of Atherosclerosis (MESA)[8], which is a multi-omics pilot study of the NHLBI Trans-Omics for Precision Medicine (TOPMed) consortium. It comprises gene expressions obtained from whole genome sequencing, as well as transcripts per million (TPM) values derived from RNA-Seq. The samples originate from four distinct populations: African American (AFA, *n* = 381), East Asian (ASN, *n* = 106), European (EUR, *n* = 538), and Hispanic/Latino (HIS, *n* = 292). Besides, our study used GWAS summary statistics from samples of East Asian ancestry as our target cohort for Systemic lupus erythematosus (SLE) and five genetically correlated autoimmune diseases (Sjogren’s syndrome, type 1 diabetes, ulcerative colitis, Hashimoto’s disease and Graves’ disease) in [9].

Because our exposure of interest is gene expression, we compare our method transMR to the popular MR framework that integrates gene expression study and GWAS, GSMR[5]. In detail, for cis-SNPs after LD-pruning (r^2^ < 0.3), GSMR exclude possible pleiotropic IVs by the HEIDI-Outlier test. While GSMR estimates the causal effect with IVW estimators based on remaining SNPs, we extend this framework with transMR, which models trans-ancestry eQTL effects and use them to estimate and refine the effects in the target cohort, then conduct HEIDI-Outlier test to remove pleiotropic IVs. Also, we extended a robust pleiotropy method for multivariable Mendelian randomization, MVMR-robost[10], in order to explore possible patterns of genes regulating each other’s expression levels. Our method transMR managed to identify more significant causal genes to the autoimmune diseases we considered comparing to the traditional IVW method, and the numbers of significant genes after the Benjamini-Hochberg correction are shown in Table (3); Most of these genes are novel findings for East Asian population (see Supplemental Materials for the full list of genes).

**Table 3.** The numbers of significant causal genes for SLE after the Benjamini-Hochberg correction from IVW and transMR. The novel genes from transMR methods are defined as causal genes that are not identified by IVW. P-values are ACAT aggregated and BH adjusted (See Materials and Methods for details).

| SLE study | Method | Number of Significant Genes | Top 5 Novel genes (ordered by adjusted p-values) | Effect | S.E. | Adjusted P-value |
| --- | --- | --- | --- | --- | --- | --- |
| 1 | IVW | 66 | - |  |  |  |
| | transMR | 68 | *STK19* | 0.369 | 0.0490 | $1.58 \times 10^{-13}$ |
| | | | *HLA-F* | -0.263 | 0.0595 | $2.43 \times 10^{-05}$ |
| | | | *HLA-K* | 0.146 | 0.0328 | $2.51 \times 10^{-05}$ |
| | | | *HLA-DPB2* | -0.164 | 0.0449 | $6.06 \times 10^{-04}$ |
| | | | *PITPNB* | -0.251 | 0.0707 | $1.04 \times 10^{-03}$ |
| | transMR-c | 64 | *Lnc-HLA-B-2* | -0.292 | 0.0377 | $3.23 \times 10^{-14}$ |
| | | | *STK19* | 1.10 | 0.186 | $1.24 \times 10^{-08}$ |
| | | | *HLA-DQA1* | -0.433 | 0.0754 | $2.69 \times 10^{-08}$ |
| | | | *HLA-K* | 0.108 | 0.0235 | $8.18 \times 10^{-06}$ |
| | | | *HLA-F* | -0.244 | 0.0536 | $1.02 \times 10^{-05}$ |
| | transMR-cBIC | 63 | *Lnc-HLA-B-2* | -0.297 | 0.0384 | $3.08 \times 10^{-14}$ |
| | | | *STK19* | 1.10 | 0.186 | $1.24 \times 10^{-08}$ |
| | | | *HLA-DQA1* | -0.493 | 0.0863 | $3.31 \times 10^{-08}$ |
| | | | *HLA-K* | 0.117 | 0.0228 | $9.26 \times 10^{-07}$ |
| | | | *HLA-F* | -0.263 | 0.0595 | $2.82 \times 10^{-05}$ |
| 2 | IVW | 15 | - |  |  |  |
| | transMR | 18 | *ZNF577* | -0.547 | 0.138 | $1.71 \times 10^{-04}$ |
| | | | *HCP5* | -0.639 | 0.168 | $3.97 \times 10^{-04}$ |
| | | | *KLF2* | 0.917 | 0.255 | $6.48 \times 10^{-04}$ |
| | | | *CLEC12A* | -0.190 | 0.0568 | $1.02 \times 10^{-03}$ |
| | transMR-c | 14 | *ZNF577* | -0.576 | 0.143 | $6.35 \times 10^{-05}$ |
| | | | *HCP5* | -0.639 | 0.168 | $4.14 \times 10^{-04}$ |
| | | | *RSPRY1* | -0.658 | 0.177 | $5.95 \times 10^{-04}$ |
| | transMR-cBIC | 15 | *ZNF577* | -0.577 | 0.143 | $6.36 \times 10^{-05}$ |
| | | | *HCP5* | -0.639 | 0.168 | $4.12 \times 10^{-04}$ |
| | | | *RSPRY1* | -0.654 | 0.175 | $5.59 \times 10^{-04}$ |
| | | | *HCG27* | 0.456 | 0.125 | $7.89 \times 10^{-04}$ |
| 3 | IVW | 3 | - |  |  |  |
| | transMR | 5 | *MMP25-AS1* | -1.05 | 0.270 | $2.26 \times 10^{-04}$ |
| | | | *STK32C* | 0.237 | 0.0616 | $2.29 \times 10^{-04}$ |
|  | transMR-c | 2 | - |  |  |  |
|  | transMR-cBIC | 3 | - |  |  |  |

First, transMR identified more causal genes from the HLA region. A strong association between the HLA region and autoimmune disease has been recognized for decades[11]. TransMR was able to pick up the well-known genetic effects such as *HLA-DQA1* on Type I diabetes (T1D) and *HLA-B* on Grave’s disease (GD)[12]. Existing evidence from studies on the East Asian population supports the causal effects of genes such as *HLA-C* for Ulcerative colitis (UC)[13] and *MICA* for Hashimoto’s disease (HD)[14]. While previous works have limited their recognition of associations to HLA classes I and II, we noticed that transMR marked some causal genes from HLA class III, such as *HLA-F* on SLE, *HLA-H* on HD, and *C4B* on GD.

Besides, several genes were identified as causal for more than one autoimmune disease. Krüppel-like factor 2 (*KLF2*) is involved in regulatory pathways of regulatory T cells and apoptotic cell clearance, which are related to autoimmunity prevention[15]. *KLF2* had a significant causal effect on GD and SLE, and the latter relationship agrees with a GWA study on the Northern Chinese population[16]. Other causal genes in common are *MORN3* (T1D and HD) and *RPL23AP1* (UC and HD).

Finally, for SLE specifically, transMR managed to make a causal conclusion for genes that were previously reported to be associated with the disease besides the *KLF2* mentioned before, naming *MBP*[17], *STK19*[18], and *NEAT1*[19]. Genes such as *PSTPIP1* and *CLEC12A* possibly are on the pathological pathways of SLE, as the former was reported to control immune synapse stability in human T cells[20], and the latter was reported to be downregulated by SLE immune complexes in normal human peripheral blood mononuclear cells[21]. Trans-ancestry extended MVMR analysis was able to identify genes *PBX2, TNXB* and *HLA-DRB1* with direct causal effects on the disease trait, compared to the default MVMR analysis. *PBX2* was reported to be aberrantly methylated in T cells, B cells, and/or monocytes in SLE patients[22]. *TNXB* was reported to be a candidate gene susceptible to SLE in the Japanese population[23]. *HLA-DRB1*, belonging to HLA Class II, has multiple detected risk alleles for SLE in the East Asian population[24] [25] [26].

We conducted the same set of analyses for the population of European ancestry based on the summary statistics for SLE. TransMR managed to identify a comparable number of causal genes to IVW (188 vs 192) for all genes available for MR after LD pruning and HEIDI-Outlier test. Then we identify potentially East Asian-specific causal genes including *PSTPIP1* and *MBP* (see Supplemental Materials for the full list of genes). Although GWA studies have reported risk loci and ancestry-specific risk loci for SLE, limited studies tested for transcriptome-wide association to make inferences on the gene expression level for the East Asian population (see [27]). Our findings shed new insights into genes of risk for SLE.

Finally, we conducted computational drug repurposing analysis using the Connectivity Map (CMap) for Systemic Lupus Erythematosus (SLE) and Graves’ disease with significant causal genes and identified several therapeutic targets. For SLE, current treatments and medications include anti-inflammatory and immunosuppressive drugs such as antimalarial drugs and glucocorticoids[28]. Anti-inflammatory drugs are beneficial in treating autoimmune diseases like Systemic Lupus Erythematosus (SLE) because they help mitigate the overactive immune response characteristic of these diseases. SLE involves chronic inflammation driven by autoantibodies targeting various cellular components, leading to systemic tissue damage, particularly in organs like the kidneys, skin, and joints. Anti-inflammatory drugs, such as glucocorticoids, reduce cytokine production, diminish immune cell recruitment to inflammation sites, and lower overall immune system activity, thereby reducing tissue damage and alleviating symptoms. Antimalarial drugs, like hydroxychloroquine and, as identified, Artesunate, offer specific benefits for autoimmune conditions due to their immunomodulatory effects beyond their traditional use. They inhibit certain pathways, such as Toll-like receptor (TLR) signaling, which plays a role in immune activation and the production of inflammatory cytokines. By interfering with these pathways, antimalarials help regulate the immune system, reducing inflammation and autoimmune activity. Further, inhibitors like IKK-2 target critical nodes in inflammatory pathways, such as the NF-κB pathway, which is pivotal for immune response regulation. NF-κB activation leads to the transcription of various pro-inflammatory genes; thus, blocking IKK-2 can dampen the downstream inflammatory response. This mechanism is especially relevant in SLE, where persistent NF-κB activity contributes to ongoing immune activation. Similarly, kinase inhibitors (e.g., H-7, Tyrphostin-AG-835), including MEK1/2 inhibitors, offer therapeutic potential by intervening in specific signaling pathways essential for immune cell activation, thereby providing a targeted approach to modulate the immune response without broadly suppressing the immune system. Besides, Janus kinase inhibitors are of interest for treating SLE[37] and we identified BRD-K06817181. For Graves’s disease, immunosuppressors such as Prednisolone, Tofacitinib, and Triptolide were identified.

## Discussion

We introduced transMR, a flexible method for inferring causal effects between pairs of traits based on summary statistics from trans-ancestry GWAS. TransMR uses a meta-regression model for SNP-exposure effects that enables aggregating summary statistics from ancestry-stratified GWAS on the same phenotype as more and more multi-ancestry genotype databases occur, which also enables winner’s curse bias correction. We showed by both simulation and real data analysis causal effect estimates from our proposed framework have smaller MSE, better power and controlled type-I error, which are especially crucial when the sample sizes of non-European cohorts are small. By further extending our framework to some existing MR methods based on summary level data (GSMR and MVMR-robust), more robust MR analysis can be conducted for the existence of pleiotropy. As our method is general, transMR can be further extended to accommodate other MR methods, e.g., MRPRESSO[29] and MRAID[30] for robust MR analysis under other assumptions, such as the InSIDE assumption[28].

Several things need to be noted when applying transMR methods to simulated or real data: (1) we used ridge to avoid overfitting as a technical consideration, while the performance was not sensitive towards the choice of λ in our simulation; (2) We need to emphasize that although it has been shown that integrating multi-ancestry information improves the power, it is still important to exclude other confounders if possible when modelling genetic effects across ancestries, since PCs of allele frequency matrix capture population stratification only. The ideal scenario to apply transMR is when summary statistics of association from large-scale, trans-ancestry GWAS are available; (3) while transMR is easier to implement with existing R function *glmnet*, transMR-c methods require customized likelihood function and extra optimization steps, which may make the method less robust, as shown in our simulation.

We also would like to mention that we show through simulation (Supplemental Materials) that recruiting IVs from other ancestry doesn’t seem to have a consistent improvement in the estimation or the power a lot compared to using IVs from identical ancestry only in transMR. The idea of looking for possible IVs from ancestries originated from the spirit of borrowing as much useful information as possible, although it was not reported that the number of IVs influences the performance of IVW estimators. Our work took the common statistical approach of selecting genetic variants, that is, to include all variants associated with the exposure at a fixed level of statistical significance (when not considering pleiotropy), hence there will always be a trade-off between re-picking up a missed IV in the target ancestry and selecting a relatively weak IV which causes bias.

We illustrate that transMR shines in discovering causal genes for autoimmune disease traits for the underrepresented East Asian population in our application since transMR identified more novel causal genes within the HLA region, which serve as positive controls, and outside the HLA region but are reported to function in immune initiation and response. On the other hand, when the target population has a large enough sample size, transMR may suffer from missing data or non-informative external data, hence would not notably improve the power, as proved when we implemented transMR in European samples.

## Methods and Materials

### Two-sample Mendelian Randomization

A classic two-sample Mendelian Randomization analysis framework is described as follows. Two-sample MR considers two separate studies in the setting of GWAS: the GWAS that measures both the exposure and the genotype data on *n*_1_ samples; and the GWAS that measures both the outcome variable of interest and the genotype data on *n*_2_ samples, without sample overlapping. If the selection of genetic instruments is done on another GWAS for exposure, where the cohort has no overlapping individuals with the exposure and disease cohorts, we call this a three-sample design. Denote the genotype matrix for the first study as ***Z*** and the *n*_1_-vector of exposure as ***x***. Denote the genotype matrix for the second study as 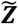 and the *n*_2_-vector of the outcome as ***y***. Assume the following linear structural equation model[31].

In the first step, for the cohort that measures the genetic instrument and the exposure, we model the association as:

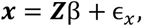

We then impute the exposure based on the effect estimates from step 1, estimate the expected exposure values and test for the association between the estimated exposure and the outcome, i.e.,

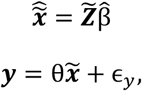

where 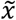 is unobserved for the second sample, and 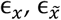 and ϵ_*y*_ denote the random error term for each equation respectively. The goal of MR methods is to make inferences on the causal effect θ, which relies on each of the selected IVs satisfying the IV assumptions mentioned before. Depending on data availability, one can use two-stage least squares (2SLS) methods for individual-level data or inverse-variance weighting (IVW) methods for summary-level data to make inferences on θ. If all IVs are perfectly uncorrelated, the estimate for θ from these two methods are similarly efficient[32]. Our method focuses on summarized data because GWAS summary statistics are more accessible for studies involving samples of diverse ancestries.

In summary-level data, for one genetic variant 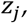 only 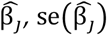 and corresponding allele frequency for *x*_*j*_ are given. Denote the estimated association and corresponding standard error of *j*_*th*_ genetic variant and *Y* in GWAS summary statistics as 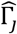 and 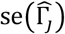. To serve as a valid instrument, the genetic variant needs to satisfy the following 3 core assumptions:

- Relevance: The IV is associated with exposure.
- Exchangeability: The IV is not associated with confounders of exposure-outcome association.
- Exclusive: The IV is only associated with the outcome through exposure.

The Wald ratio estimator for the causal effect θ from the *j*_*th*_ instrument is

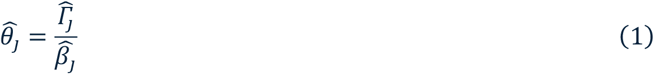

with first-order approximate standard error

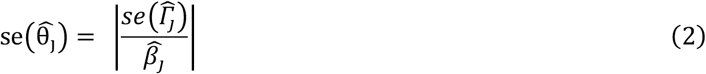

If all instruments are independent, valid instruments, we can obtain a consistent estimator for θ by the usual inverse-variance weighted (IVW) regression:

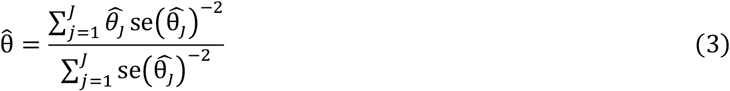

and its standard error is:

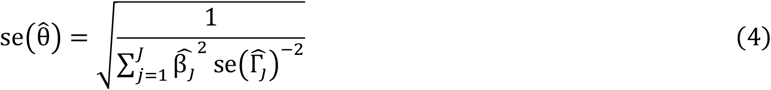

### Trans-ancestry two-sample Mendelian Randomization

To accommodate the situation when multi-ancestry exposure cohorts are available, we assume *K* cohorts are available and index different ancestries by *k, k* = 1, …, *K*, and model each β_*j*_ with a fixed effect of the ancestry captured by the allele frequency (AF) principal components. Denote the (*K* + 1) × *J* allele frequency matrix of genetic variants shared in all cohorts, along with AF for the same set of variants from the reference panel (e.g., 1000 Genomes Project) as *F* and its principal components (PCs) as ***W***_1_, …, ***W***_*K*_, assuming *J* ≥ (*K* + 1). For the *j*_*th*_ instrument, we model its effect on exposure as

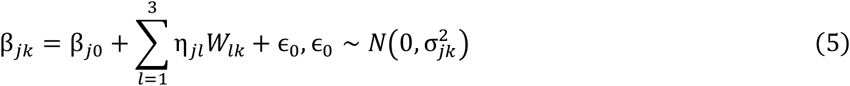

which is a weighted linear regression model with larger weights assigned to variant-exposure GWAS cohorts with smaller se 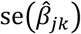. Usually, this weight is set to 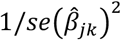. Each instrument has an individual model, and these models are not related to each other. For instruments unaffected by ancestry, coefficient estimates are expected to remain close to zero. To prevent overfitting in these estimates, we introduce a ridge penalty with a small regularization parameter λ ∈ {0.01,0.5,1} penalizing for large values of η_*j*_. The ridge penalty modifies the standard loss function by adding a regularization term, expressed as follows:

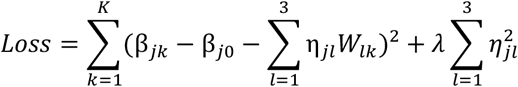

The winner’s curse bias correction is done by replacing the contribution from the target population, say cohort k, with a log-conditional probability in the ridge penalized log-likelihood for (5):

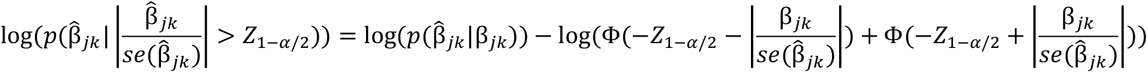

while 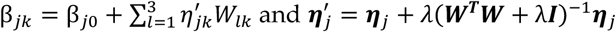 which accounts for the bias brought by the ridge penalty.

Then we make a prediction of β_*j,K*+1_ based on the previous model with ***W***_*K*+1_, and denote the predicted values as 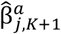. The estimator, standard error for θ_*j*_ and furthermore, θ can be obtained by simply replacing 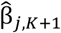 with 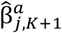 when using equation (1), (2) and (3).

We also consider a 3-sample design as a gold standard to compare with TransMR. Specifically, if another independent exposure sample is available, we can also conduct a 3-sample design, where we use one exposure sample to select the instrument variable and another sample to estimate the genetic effect on the exposure, which can overcome the inflation caused by the winner’s curse. Yet, the availability of an independent GWAS on the exposure is questionable, particularly for non-European populations. So, the consideration of a 3-sample design is theoretical, which can be used as an upper bound for power.

### Extension of GSMR and MVMR

Now we show how to extend existing popular MR methods, in particular, GSMR and MVMR based on our method, so they can be implemented in our data application of identifying causal genes for autoimmune disease traits from samples of diverse ancestries.

Both SMR[33] and GSMR have been applied to perform integrative analysis of gene expression study and GWAS. Specifically, SMR selects one SNP that has the smallest p-value of the marginal association test in the cis-region of the gene to serve as the IV. Afterwards, SMR uses the standard MR ratio method to estimate the causal effect. Different from SMR, which uses only one IV, GSMR selects multiple independent SNPs in the cis-region to serve as IVs. In particular, after LD-pruning, GSMR tests for and rules out possible pleiotropic variants through the HEIDI-Outlier test, estimates the causal effect of each IV in turn using the standard MR ratio method, and combines these causal effect estimates with the standard IVW approach. Next, we illustrate how to conduct the HEIDI-Outlier test after obtaining 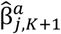.

Denote the weighting matrix for model (5) as ***M***. The variance of estimation for η_*j*_ from the regular weighted linear regression forms as

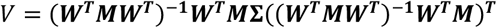

where **Σ** = Var(ϵ). The ridge estimator for η_*j*_, can be represented as a linear transformation of the weighted least square estimator:

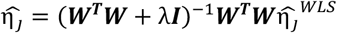

and we have

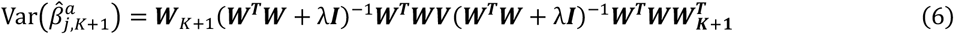

The basic idea of HEIDI-Outlier is to test where there is a significant difference between the estimation from any instrument, 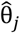, and estimation from a target SNP that shows a strong association with the exposure, 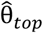. The difference 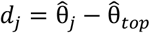 has a variance 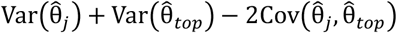. Use the χ^2^-statistic 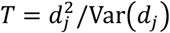 and remove the SNPs with p-value < 0.01.

Multivariable MR (MVMR) is an extension to MR that uses genetic variants associated with multiple, potentially related exposures to estimate the effect of each exposure on a single outcome. This allows for mediation analysis and therefore can be used to estimate gene regulatory effects[34]. Based on summary-level data only, this is done by fitting the following weighted regression model:

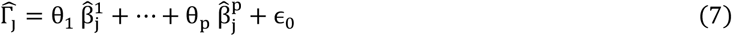

While univariable MR estimates the total effect of the exposure on the outcome, MVMR estimates the marginal effect of the exposure on the outcome conditional on the mediators. The difference between these estimates (i.e., “difference in effects”) will then give the mediated effects. In order to overcome pleiotropic effects, MVMR-Robust described in [10] uses robust regression methods MM-estimation. In a multi-ancestry setting, we simply plug in the corresponding 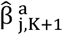 (and variance estimated by equation (6) into (7).

### Simulation Studies

We generated individual-level data of independent SNPs (***Z***), generic continuous exposure (**Σ**), and outcome (*Y*) for six cohorts (5 with exposure measured, 1 with outcome) of different ancestry of European (EUR), African (AFR), East Asian (ASN), Hispanic (HIS), Brazilian (BRZ), and European (EUR), with independent samples. For the scenario of weak instruments, we set the sample size to 15k to ensure basic power. For the scenario of the underrepresented population, we set the cohorts as (AFR, EUR, ASN, ASN, AFR, AFR), and sample sizes for EUR ancestry as 20000, while ASN and AFR are of 10% and 5%. Otherwise, all cohorts have the same sample size equal to 5000.

In each exposure cohort, the variables are generated according to the following structural equations:

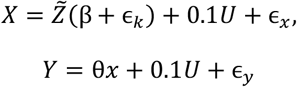

where β_*j*_’s ∼*N*0.1, 0.05^2^). For the scenario of weak instruments, we set β_*j*_ ∼ *N*(0.08,0.05^2^) and β_*j*_ ∼(0.05,0.05^2^). We use the reference population-stratified allele frequencies reported in [35] to generate genotypes. Out of 3076 SNPs, we randomly selected 25 as the pool of causal variants for *X* and 20 out of the pool for each ancestry. SNPs *N* = (*N*_1_, …, *N*_20_) denotes the SNP counts (0,1,2) for each causal variant and were generated according to the reference AF, and 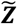 is the column-wise standardized genotype matrix. Confounder, *U*, is generated by a normal distribution with fixed pre-specified mean and variance (−0.5,4). The random error terms ϵ_*x*_, ϵ_*y*_ follow the standard normal distribution. The heterogeneity in β’s is generated by ϵ_k_ ∼*N*(0, (0.02φ_*k*_)^2^), which denotes the scale parameter of *k*_*th*_ ancestry among EUR, AFR, ASN, HIS and BRZ and is equal to 0, φ, 2φ,0.5φ,0.5φ respectively.

We ran a single-trait association test on each cohort using the package GENESIS[36] and select SNPs with p-values smaller than 1 × 10^−3^ which is equivalent to IV strength defined by the approximated F-statistic 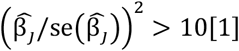 For multi-ancestry exposure cohorts, we select IVs for the ancestry that matches the outcome study and use them with TransMR for analysis. For example, if the outcome cohort is East Asian, we will use the exposure cohorts with East Asian ancestry to select our IVs.

### Application

We conducted eQTL analysis on the MESA RNASeq data obtained from [37] with the package MatrixEQTL[38]. For the genotype data excluding chromosome X, we performed quality control (QC) that removed SNPs that are INDELs, multiallelic, and ambiguous-strand (A/T, T/A, C/G, G/C) and removed the remaining variants with minor allele frequencies (MAFs) < 0.01 and Hardy-Weinberg Equilibrium (HWE) *p* < 1 × 10^−6^. We filtered for significant cis-eQTLs with p-value < 1e-3 in the ASN population, conducted LD-pruning, and extracted the same set of eQTLs from other ancestries. After filtering for SNPs available in GWAS summary statistics, significant eQTLs and reference panel, and harmonization of alleles, we perform eQTL analysis for more than 7,000 genes. For ridge regression on β_*jk*_, we use the first 2 PCs of the AF matrix since there are only four exposure cohorts.

To ensure the power after the meta-regression model of eQTL effects, we use 0 PC (i.e., the fixed-effect model), 1 PC, and 2 PCs in the meta-regression step, and based on predicted eQTLs effects, we obtain estimations for causal effect respectively. Then we use the Cauchy combination test [39] to combine individual p-values from 3 pairs of estimations. We applied Benjamini-Hochberg correction on the p-values for each set of results.

## Acknowledgement

MESA-TOPMed/MESA Study Acknowledgement

MESA and the MESA SHARe project are conducted and supported by the National Heart, Lung, and Blood Institute (NHLBI) in collaboration with MESA investigators. Support for MESA is provided by contracts HHSN268201500003I, N01-HC-95159, N01-HC-95160, N01-HC-95161, N01-HC-95162, N01-HC- 95163, N01-HC-95164, N01-HC-95165, N01-HC-95166, N01-HC-95167, N01-HC-95168, N01-HC-95169, UL1-TR-000040, UL1-TR-001079, UL1-TR-001420. MESA Family is conducted and supported by the National Heart, Lung, and Blood Institute (NHLBI) in collaboration with MESA investigators. Support is provided by grants and contracts R01HL071051, R01HL071205, R01HL071250, R01HL071251, R01HL071258, R01HL071259, and by the National Center for Research Resources, Grant UL1RR033176. This study was supported in part by the National Center for Advancing Translational Sciences, CTSI grant UL1TR001881, and the National Institute of Diabetes and Digestive and Kidney Disease Diabetes Research Center (DRC) grant DK063491 to the Southern California Diabetes Endocrinology Research Center. The TOPMed MESA Multi-Omics project was conducted by the University of Washington and LABioMed (HHSN2682015000031/HHSN26800004). Molecular data for the Trans-Omics in Precision Medicine (TOPMed) program was supported by the National Heart, Lung and Blood Institute (NHLBI). RNA-seq for the NHLBI TOPMed: Multi-Ethnic Study of Atherosclerosis (MESA)” (phs001416.v1.p1) was performed at the Northwest Genomics Center (HHSN268201600032I) and at the Broad Institute Genomics Platform (HHSN268201600034I). Core support including centralized genomic read mapping and genotype calling, along with variant quality metrics and filtering were provided by the TOPMed Informatics Research Center (3R01HL-117626-02S1; contract HHSN268201800002I). Core support including phenotype harmonization, data management, sample-identity QC, and general program coordination were provided by the TOPMed Data Coordinating Center (R01HL-120393; U01HL-120393; contract HHSN268201800001I). We gratefully acknowledge the studies and participants who provided biological samples and data for TOPMed.

Whole genome sequencing (WGS) for the Trans-Omics in Precision Medicine (TOPMed) program was supported by the National Heart, Lung and Blood Institute (NHLBI). WGS for “NHLBI TOPMed: Multi-Ethnic Study of Atherosclerosis (MESA)” (phs001416.v3.p1) was performed at the Broad Institute of MIT and Harvard (3U54HG003067-13S1). Centralized read mapping and genotype calling, along with variant quality metrics and filtering were provided by the TOPMed Informatics Research Center (3R01HL- 117626-02S1). Phenotype harmonization, data management, sample-identity QC, and general study coordination, were provided by the TOPMed Data Coordinating Center (3R01HL-120393-02S1), and TOPMed MESA Multi-Omics (HHSN2682015000031/HSN26800004). The MESA projects are conducted and supported by the National Heart, Lung, and Blood Institute (NHLBI) in collaboration with MESA investigators. Support for the Multi-Ethnic Study of Atherosclerosis (MESA) projects are conducted and supported by the National Heart, Lung, and Blood Institute (NHLBI) in collaboration with MESA investigators. Support for MESA is provided by contracts 75N92020D00001, HHSN268201500003I, N01- HC-95159, 75N92020D00005, N01-HC-95160, 75N92020D00002, N01-HC-95161, 75N92020D00003, N01- HC-95162, 75N92020D00006, N01-HC-95163, 75N92020D00004, N01-HC-95164, 75N92020D00007, N01- HC-95165, N01-HC-95166, N01-HC-95167, N01-HC-95168, N01-HC-95169, UL1-TR-000040, UL1-TR- 001079, UL1-TR-001420, UL1TR001881, DK063491, and R01HL105756. The authors thank the other investigators, the staff, and the participants of the MESA study for their valuable contributions. A full list of participating MESA investigators and institutes can be found at http://www.mesa-nhlbi.org

## Supplemental Materials for “Trans-ancestry Mendelian Randomization Discovers Novel Causal Genes for Autoimmune Disease Traits”

**Table S1.** Top 5 novel significant causal genes and number of significant causal genes inferred by transMR methods for autoimmune diseases of Systemic lupus erythematosus, Sjogren’s syndrome, type 1 diabetes, ulcerative colitis, Hashimoto’s disease and Graves’ disease.

| AiD trait | Method | Top 5 Novel genes<br>(ordered by adjusted p-values) | Number of<br>Significant Genes |
| --- | --- | --- | --- |
| SLE-1 | IVW | - | 66 |
|  | transMR | *STK19, HLA-F, HLA-K, HLA-DPB2, PITPNB* | 68 |
|  | transMR-c | *Lnc-HLA-B-2, STK19, HLA-DQA1, HLA-K, HLA-F* | 64 |
|  | transMR-cBIC | *Lnc-HLA-B-2, STK19, HLA-DQA1, HLA-K, HLA-F* | 63 |
| SLE-2 | IVW | - | 15 |
|  | transMR | *ZNF577, HCP5, KLF2, CLEC12A* | 18 |
|  | transMR-c | *ZNF577, HCP5, RSPRY1* | 14 |
|  | transMR-cBIC | *ZNF577, HCP5, RSPRY1, HCG27* | 15 |
| SLE-3 | IVW | - | 3 |
|  | transMR | *MMP25-AS1, STK32C* | 5 |
|  | transMR-c | - | 2 |
|  | transMR-cBIC | - | 3 |
| T1D | IVW | - | 11 |
|  | transMR | *HLA-DQA1, Lnc-TUBB2A-7, HLA-DRB5, MORN3, DPY19L1* | 17 |
|  | transMR-c | *HLA-DQA2, HLA-DRB6, HTATSF1P2, DLG5, HLA-DQA1* | 14 |
|  | transMR-cBIC | *HLA-DQA1, Lnc-TUBB2A-7, MORN3, DPY19L1* | 13 |
| Hashimoto | IVW | - | 26 |
|  | transMR | *HLA-DRB6, MRPL43, HLA-DRB5, RPL23AP1, RPS18* | 33 |
|  | transMR-c | *HLA-DRB1, HLA-DRB6, HCG4, HLA-C, Lnc-MRPL43-2* | 31 |
|  | transMR-cBIC | *HLA-DRB6, HLA-DRB5, ATP6V1G2-DDX39B, MRPL43, TNXB* | 31 |
| Graves | IVW | - | 55 |
|  | transMR | *HLA-DRB5, HLA-B, SNHG32, C11orf21, C4B* | 57 |
|  | transMR-c | *HCG4, HLA-H, HLA-DRB5, HLA-DQA1, HLA-DQA2* | 62 |
|  | transMR-cBIC | *HLA-DRB5, C4B, C11orf21, HLA-B, GPSM3* | 62 |
| UC | IVW | - | 23 |
|  | transMR | *ATP6V1G2-DDX39B, NCR3, HLA-C, HCG27, RPL23AP1* | 28 |
|  | transMR-c | *HLA-DRB6, HLA-DQB2, ATP6V1G2-DDX39B, HLA-DQA1, MICA* | 25 |
|  | transMR-cBIC | *ATP6V1G2-DDX39B, RPL23AP1, NCR3, HLA-C, HLA-B* | 28 |
| Sjogren | IVW | - | 26 |
|  | transMR | *HLA-DQA2, TMEM44, HLA-DPB2, PANX1, FIGNL1* | 26 |
|  | transMR-c | *HLA-DQA2, HLA-DRB1, RING1, HLA-DRB6, HLA-DQA1* | 35 |
|  | transMR-cBIC | *HLA-DQA2, TMEM51, TNXB, CARS2, ZNF611* | 34 |

**Table S2.**
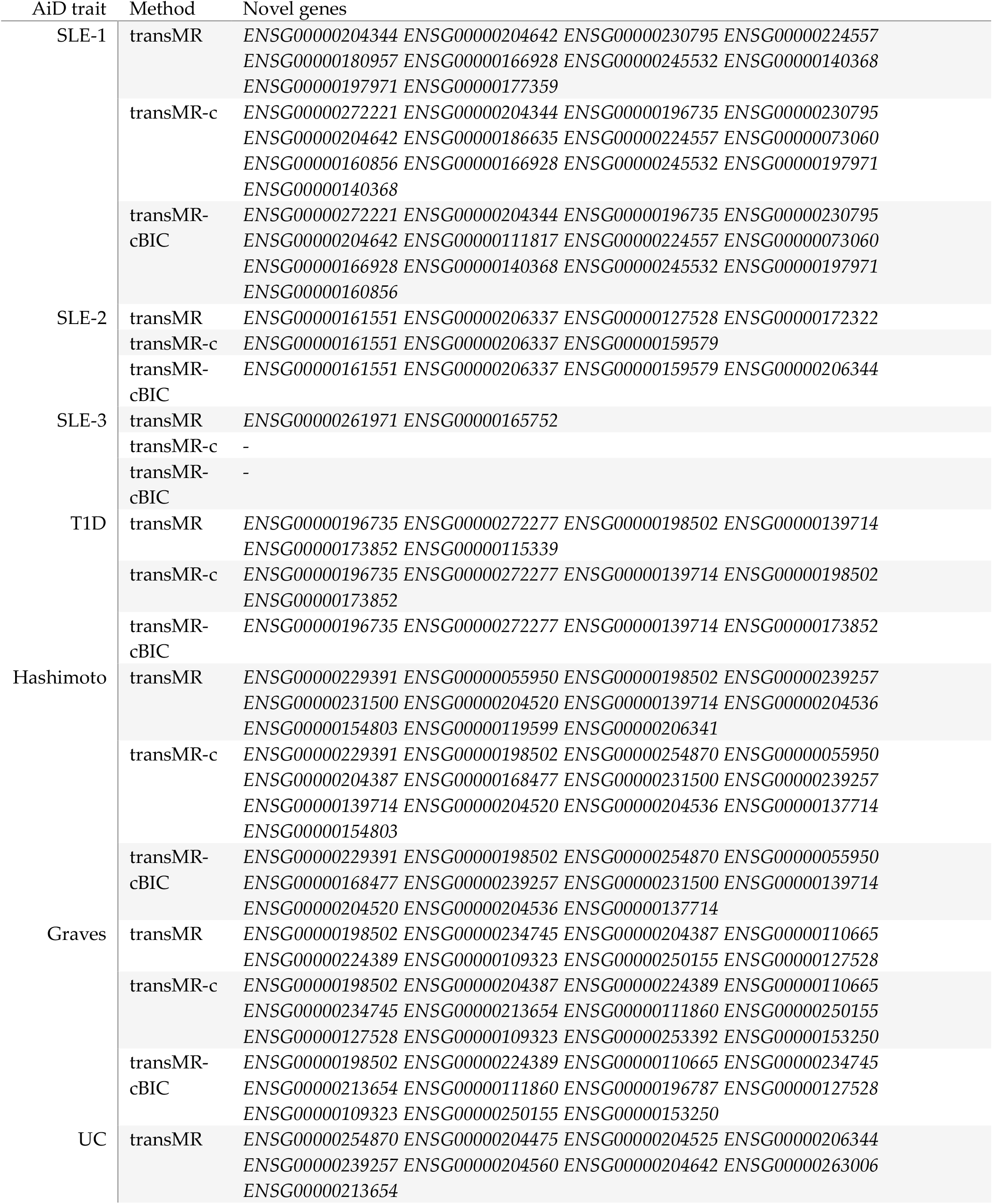
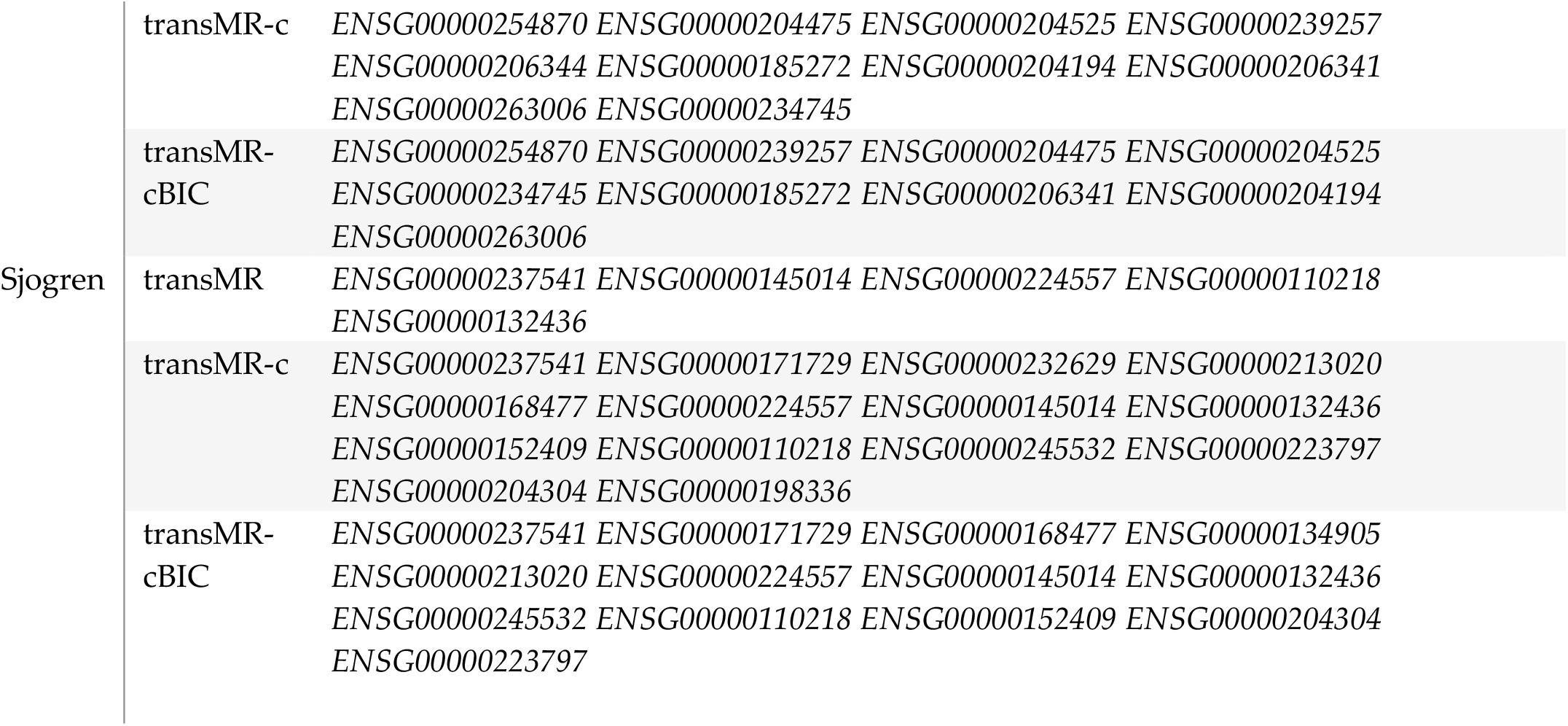
Full list of novel causal genes inferred by transMR methods for autoimmune diseases of Systemic lupus erythematosus, Sjogren’s syndrome, type 1 diabetes, ulcerative colitis, Hashimoto’s disease and Graves’ disease.

**Figure S3.**
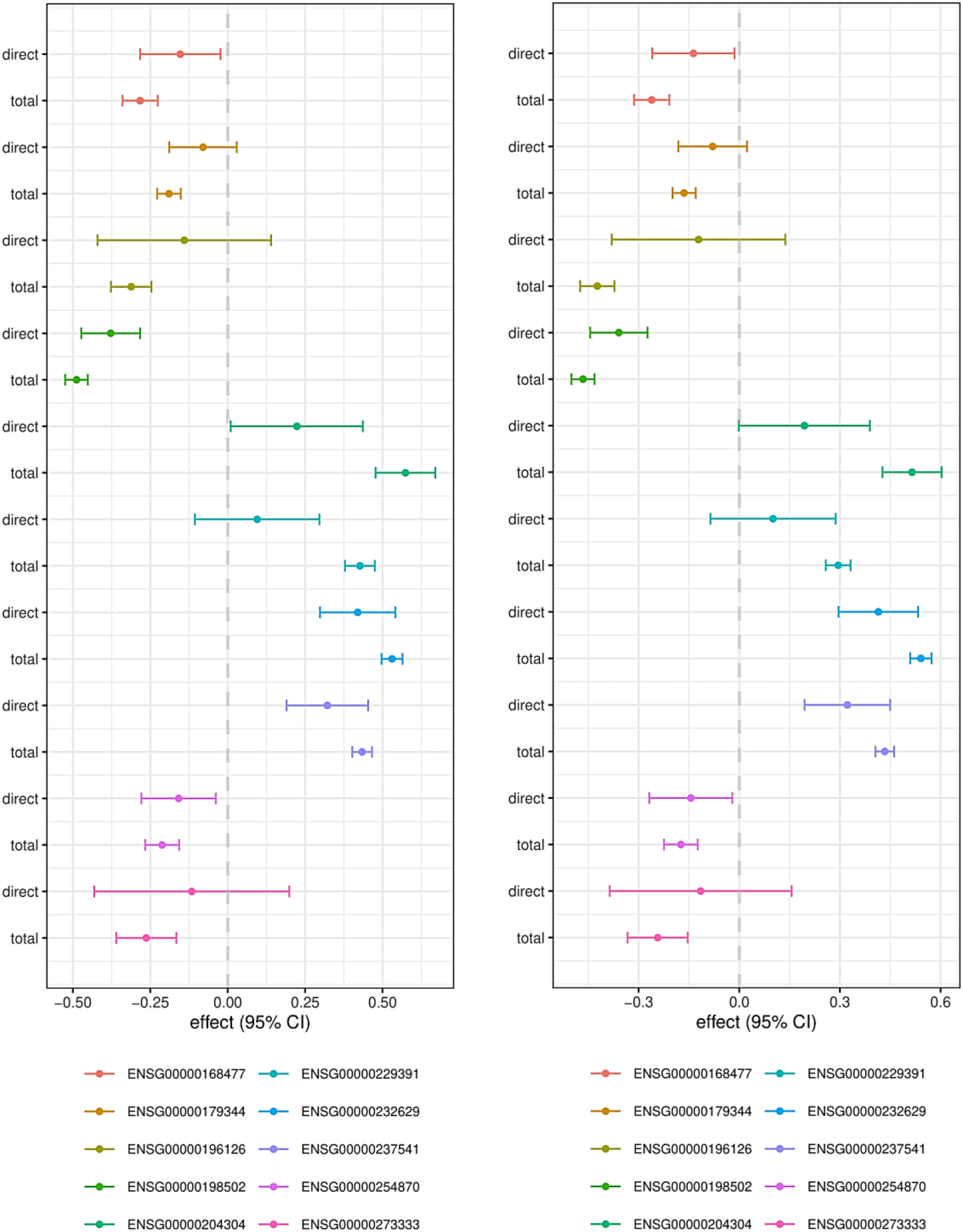
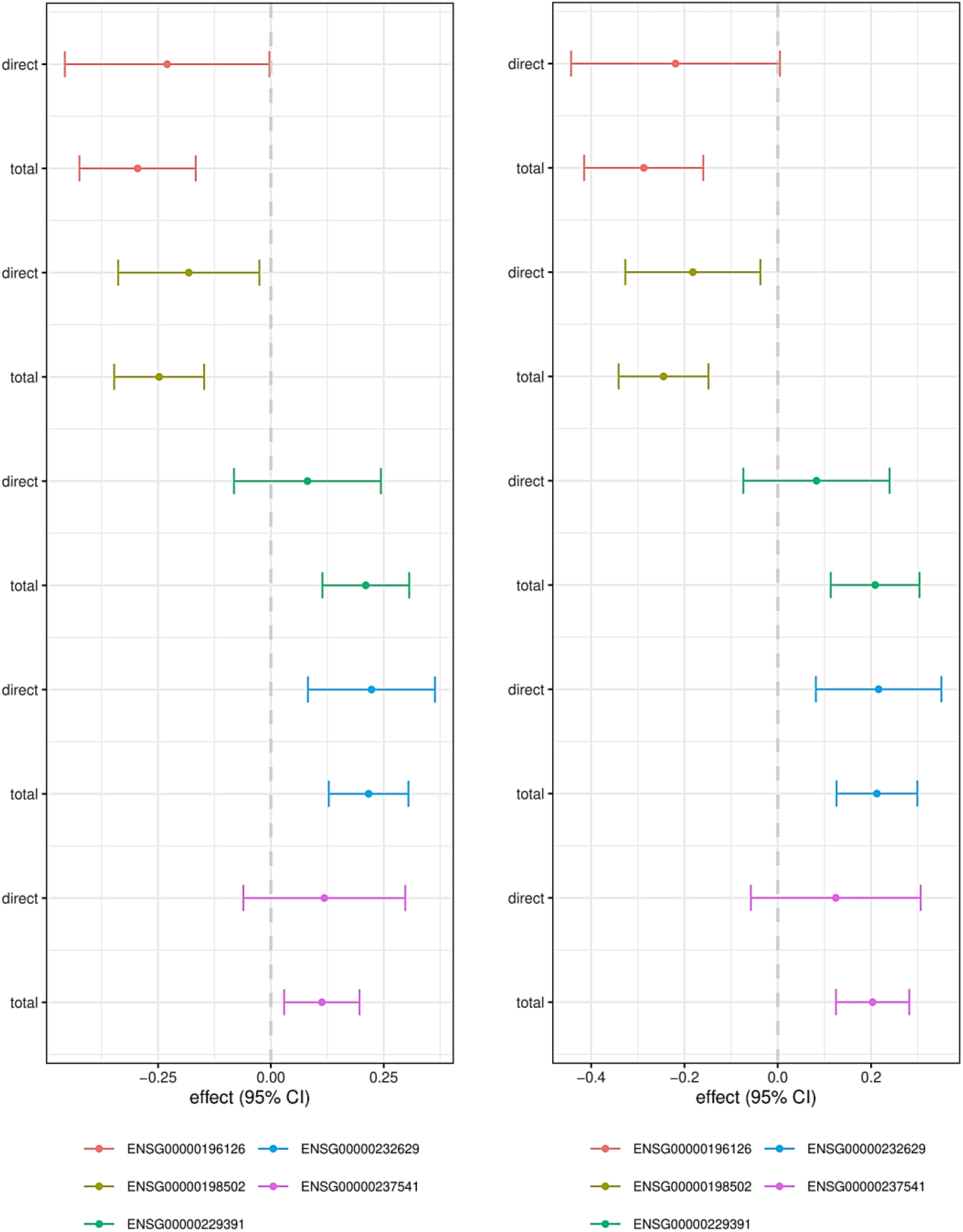
Multivariate Mendelian Randomization results for causal genes of Systemic lupus erythematosus study 1 and 2. *PBX2* and *TNXB* showed significant direct effect from SLE study 1 (up left), while *HLA-DRB1* showed significant direct effect from SLE study 2 (lower left), and the effects of *PBX2* and *HLA-DRB1* are borderline non-significant when using default MVMR-Robust method (up right, lower right).

**Table S4.** TransMR inferred causal genes for SLE in European ancestry population. Significant genes are evaluated on genes that are valid for both inference methods.

| AiD trait | Method | Top 5 Significant causal genes | Novel genes | Number of Significant Genes |
| --- | --- | --- | --- | --- |
| SLE | IVW | *ENSG00000269918*<br>*ENSG00000255310*<br>*ENSG00000154319*<br>*ENSG00000173295*<br>*ENSG00000186470* | - | 192 |
|  | transMR | *ENSG00000173295*<br>*ENSG00000269918*<br>*ENSG00000186470*<br>*ENSG00000154319*<br>*ENSG00000255310* | *ENSG00000140678*<br>*ENSG00000183386*<br>*ENSG00000212978*<br>*ENSG00000149591*<br>*ENSG00000151690*<br>*ENSG00000122223*<br>*ENSG00000101161*<br>*ENSG00000164308*<br>*ENSG00000070190*<br>*ENSG00000254859*<br>*ENSG00000280670*<br>*ENSG00000198040*<br>*ENSG00000186792*<br>*ENSG00000197208*<br>*ENSG00000237651*<br>*ENSG00000198756*<br>*ENSG00000116761*<br>*ENSG00000126790*<br>*ENSG00000196502*<br>*ENSG00000176476*<br>*ENSG00000138604*<br>*ENSG00000115364*<br>*ENSG00000139190* | 188 |

**Figure S5.**
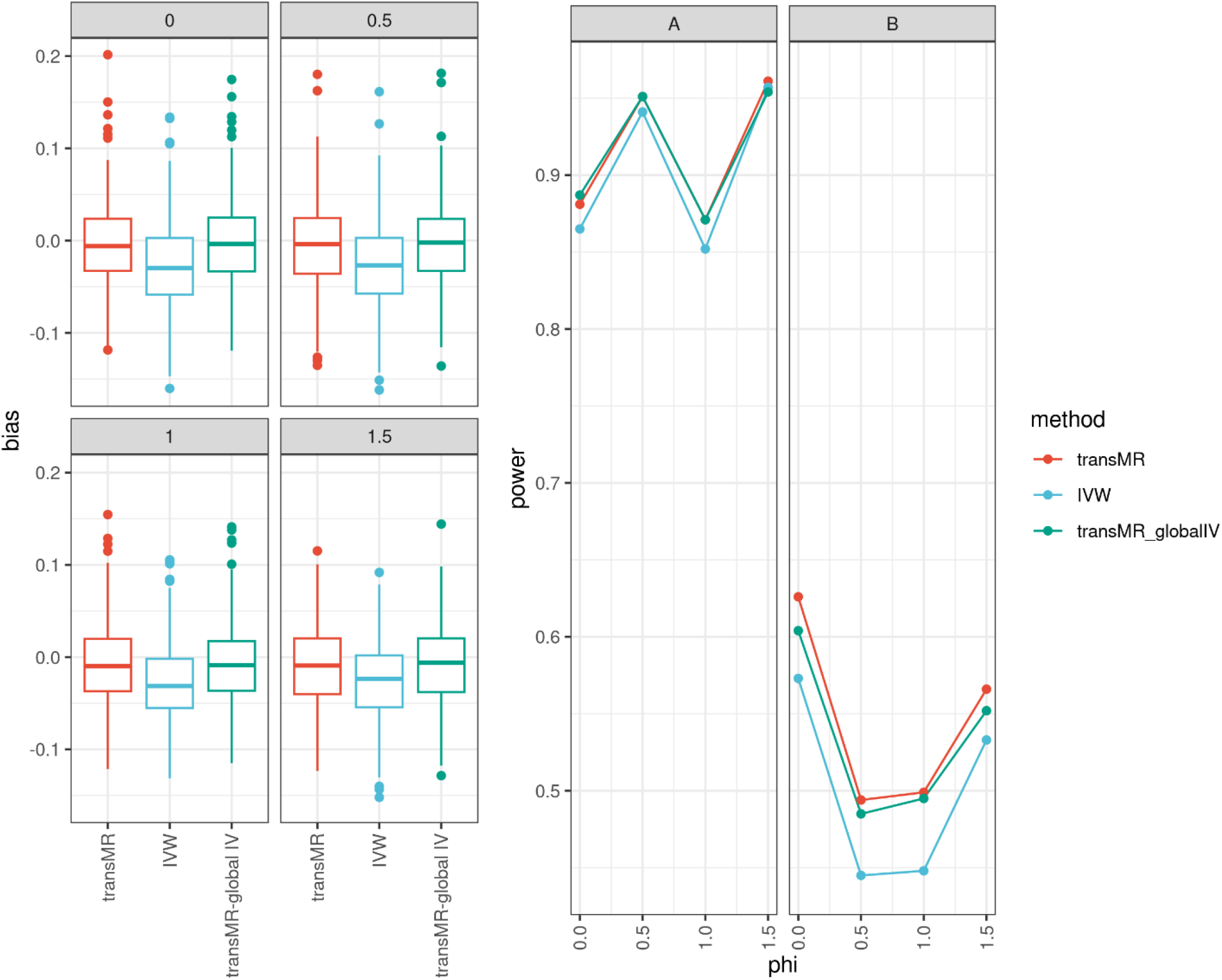
Recruiting IVs from other ancestry (labelled as “transMR-global IV”) doesn’t seem to have a consistent improvement in the estimation or the power a lot compared to using IVs from identical ancestry only in transMR. Left: bias from transMR, IVW and transMR-global IV estimators under different levels of heterogeneity in SNP effects in exposure, which is controlled by *φ*; Right: Power of transMR, IVW and transMR-global IV when the true causal effect = 0.1 with 10k samples in each cohort, and for senarios (A) the heterogeity in SNP effects is small; (B) all the IVs are weak.

## Notes

### Competing Interest Statement

The authors have declared no competing interest.

